# Predicting Distant Recurrences in Invasive Breast Carcinoma Patients Using Clinicopathological Data: A cross-institutional and AI-based study

**DOI:** 10.1101/2023.04.16.537076

**Authors:** Shrey S. Sukhadia, Kristen E. Muller, Adrienne A. Workman, Shivashankar H. Nagaraj

## Abstract

Breast cancer ranks second in the most common cancer in women worldwide with 30% of cases resulting into recurrence of the disease at distant organs post the treatment. While clinicians have utilized several clinicopathological measurements for prediction of distant recurrences in invasive breast carcinoma (IBC), none of those studies have showcased the potential of combining clinicopathological evaluations of IBC tumors pre and post therapies using machine learning (ML) or artificially intelligent (AI) models to predict the distant recurrence of the disease in respective patients. The goal of our study was to determine whether classification-based ML/AI techniques can predict distant recurrences in IBC patients using key clinicopathological measurements that includes pathological staging of tumor and surrounding lymph nodes deemed both pre- and post-neoadjuvant therapy, imaging-based therapy responses, and the status of adjuvant therapy administered to patients. We trained and tested clinicopathological ML/AI model using dataset from Duke University and validated it using external dataset from Dartmouth Hitchcock Medical Center (DHMC). Random Forest (RF) model performed best compared to C-Support Vector Classifier (SVC) and Multi-Layer Perceptron (MLP) yielding AUC ranging 0.75-1.0 (p<0.002) across both the institutions, thereby demonstrating the cross-institutional portability and validity of ML/AI models in the field of clinical research in cancer.

## Introduction

Breast cancer is the most common cancer in women worldwide, affecting one in eight women. Of those, up to 30% will eventually develop metastatic disease, which is fatal for the vast majority of patients. Standardized treatment regimens, including targeted therapies, exist for a subset of breast cancer patients; however, breast cancer is a heterogeneous disease, and long-term survival rates and prognosis varies widely even within the same histologic and molecular subtype of breast cancer. An effective and efficient system to predict recurrences and metastases in breast cancer patients is crucial to the development of personalized monitoring and treatment strategies. Neoadjuvant chemotherapy, i.e. chemotherapy that precedes surgery, has become the standard of care for a subset of breast cancer patients with the intent to downstage disease and improve survival. A patient’s response to neoadjuvant chemotherapy is evaluated by clinical and pathologic examination and characterized as a complete response (no residual tumor), partial or incomplete response (reduction in tumor), or no response (no decrease in tumor size or volume). The degree of response to neoadjuvant chemotherapy is predictive of outcome; in particular, a complete pathologic response with no residual tumor is indicative of an excellent prognosis in patients with *HER2*-positive and triple-negative tumors (Provenzano et al. 2015; von Minckwitz et al. 2012).

Currently, multigene assays such as the OncotypeDX are widely used in clinical practice to predict the distant recurrence rate and benefit from adjuvant chemotherapy for patients with estrogen receptor (ER)-positive IBC (Albain et al. 2010), but the prediction of the chemotherapy benefits vary depending on the range of OncotypeDX score obtained for patients combined with their age (Sparano et al. 2018). In addition, nomogram calculators are available that provide five and ten-year disease free probabilities for patients who receive neoadjuvant chemotherapy; however, these are limited to logistic regression and Cox proportional hazard regression models performing at an AUC less than 90% and concordance-index (C-index) of less than 80%, respectively (Boutros et al. 2015; Zhang et al. 2019). Xia et al. developed a nomogram for triple-negative breast cancer patients who underwent neoadjuvant chemotherapy using a radiomic signature derived from preoperative MRI and clinicopathological variables; however, this showed a C-index of less than 90% (Xia et al. 2021). PREDICT, a UK based prognostic model based on ER and progesterone receptor (PR) status with adjustment for prognostic factors and mode of cancer detection (i.e. symptomatic or screen-detected) predicts survival post-surgery for IBC patients, but performs at AUC less than 90% (Wishart et al. 2010).

Machine learning (ML) models have been developed to predict the risk of metastatic disease for patients with early stage breast cancer, but the patients therein lack the administration of adjuvant therapy, making such analyses restricted to a group of therapy-naïve patients (Nicolò et al. 2020). Some studies have used clinicopathologic features including serum levels of HER2 (i.e. sHER2) to predict metastases in patients; however, the low AUC (i.e. AUC<0.8) achieved for their models would make it difficult to utilize such a test clinically. Furthermore, an independent validation test was not peformed (Tseng et al. 2019). The aforementioned studies do not expose transparent reports that could aid in tracing features through each step in algorithm unlike the ML/AI operational reports yielded by ImaGene (Sukhadia et al. 2022). ImaGene automates ML/AI operations and provides users with multiple model types to choose from, such as RF, SVC, MLP, MultiTask Linear Regression, and MultiTask LASSO. The software offers various parameters to experiment through and alludes to metrics such as R-square, RMSE:Stdev ratio and AUCs, majority of which are plotted as graphs and crafted into an operational report. These reports, along with the subordinate tabular output files, aid users in tracking feature-level performances for the respective models tested (Sukhadia et al. 2022).

For breast cancer patients, predicting the risk of metastases and tailoring therapy accordingly is crucial; however, clinicians are currently restricted to predict disease progression by compiling information from clinical, pathologic, and molecular testing. The aim of our study was to evaluate if clinicopathologic measurements including pre- and post-neoadjuvant clinical and pathological staging, response to neoadjuvant therapy, and administration of adjuvant therapy could be modelled together using ML/AI techniques to predict distant recurrence flags in IBC patients at higher than 90% (aka 0.9) AUC. We focused on patients who received neoadjuvant chemotherapy and had either a partial or a complete response. A variety of classification-based ML/AI models, i.e., Random Forest, Multi-layer perceptron (aka supervised neural network) and C-Support Vector Classifier (SVC) were employed for training, testing and validation of the clinicopathologic features using ImaGene (Sukhadia et al. 2022). ImaGene yielded detailed operational reports alluding features and model-parameters used during training, testing and validation for all three models. These reports and the supporting tabular output files showcased several metrics and plots collectively, which included Cross Validation (CV) score, Grid Search score (in case grid search set to “True”), Actual-vs-Predicted-values scatter plots, Area under the Receiver Operating Curve (AUC) and its respective p-value for the prediction of distant recurrence flag. The training and testing of the models were performed using a publicly available clinical dataset from one institution, Duke University, followed by the validation of the best tested model using a similar dataset from an external institution, Dartmouth Hitchcock Medical center (DHMC). This provided us with a cross-institutionalclinicopathological AI model that could be validated further at other institutions with the eventual goal to utilize such tools in clinical practice to predict distant recurrences.

## Method

The clinicopathological data from a retrospective study of 900 invasive breast carcinoma patients at Duke University (available at The Cancer Imaging Archive(TCIA) portal) were downloaded and explored in this study (Clark et al. 2013; Saha et al. 2018). These patients received neoadjuvant therapies such as chemotherapy, radiation or endocrine hormone-based therapy, had their responses evaluated through imaging and pathological staging (both tumor and nodal), and some received adjuvant therapies. The patient entries pertaining to ungraded tumor responses from therapies or those labelled as ‘Not applicable (NA)’ for any of the aforementioned clinicopathological parameters were filtered out thereby retaining 161 patient-entries. These were split further into train and test sets at 90:10 ratio, where 90% of entries were used for training (i.e., n_train_=144) and 10% for testing (n_test_=17) of several classification based ML models such as Random Forest (RF), C-Support Vector Classification (SVC) and Supervised Neural Network (aka Multi-Layer Perceptron, i.e. MLP) to predict distant recurrence flags or labels (“Yes” or “No”). Specifically, these clinicopathological parameters consisted of the following features: a) Tumor stage or size (T0-T4) pre-neoadjuvant therapy, b) Staging of lymph nodes (N0-N3), c) Clinical response to neoadjuvant therapy evaluated using radiology images (complete or incomplete response or assessment unavailable), d) Pathologic response to neoadjuvant therapy (complete, incomplete, or DCIS-only remaining), e) Pathologic response to neoadjuvant therapy evaluated using pathology stage “T” tumor size (T0-T4), f) Pathologic response to neoadjuvant therapy evaluated using node status “N” (N0-N3), g) Status of adjuvant chemotherapy administered (yes or no), and h) Status of adjuvant anti-*Her2/Neu* therapy administered (yes or no). Patients with no response to neoadjuvant chemotherapy were excluded in this study as this patient population has a poor prognosis and higher risk of recurrence (Symmans et al. 2007). The training of the model was conducted using ImaGene software (Sukhadia et al. 2022). Out of the three models, RF performed the best with grid search activated over tree-depth hyperparameter to predict distant recurrence flags in test patients (n_test_=17) at AUC=1.0 (p<0.002 calculated over n_permut_=14 permutations of the distant recurrence flags/labels).

The experiments for various model types that were performed on the ImaGene platform (Sukhadia et al. 2022) yielded transparent AI reports and the supporting text files aiding interpretation of results leading to selection of the model eligible for validation. Validation was done through an independent set of patients from DHMC that met two criteria: a) neoadjuvant therapy administered, and b) In-house (i.e. DHMC on-site) follow-up visits post adjuvant therapy. We started with a total of 67 patients, out of which 31 patients did not meet the above two criteria leading to 36 that remained. Only two patients out of these 36 had distant recurrences, leading us to select 9 patients randomly (i.e., two with and seven without distant recurrences) to balance the dataset better to arrive at a more realistic validation-AUC.

### Model development and testing using ImaGene

The multimodal feature file consisting of the aforementioned clinicopathological features were used as “data” and a binary column containing feature file indicating whether the disease recurred at a distant site (aka distant recurrence flag, i.e. 0 for “no” and 1 for “yes”) for the respective patients was used as “label” on the ImaGene platform. ImaGene was first run through a RandomForest model-type in “Train” mode with n=161 patients, setting test size to “0.1 (aka 10% of dataset allocated to test)” which partitioned the dataset into train (n_Train_=144) and test (n_Test_=17) sets. The K-fold cross validation splitter parameter (i.e., ‘cv’) was set to “2”. Grid search was set to “True” to enable execution of grid search through the hyperparameters: max_depth = [6, 9, 10, 12, 15, 20] and cv=4. The “data” normalization method was set to “StandScaler” while the “label” normalization set to “none” due to its binary nature. Further, the absolute correlation threshold parameter pre-ML-training was set to “-1.0” to silent the filtering of features based on Pearson’s correlation co-efficient threshold, thereby considering all the clinicopathological features for ML/AI model training further. The run took two minutes to yield a report that showed performance of the model (Supplementary Report 1, Figure 1a and Table 1) which is described in Results section as well.

**Figure 1:**
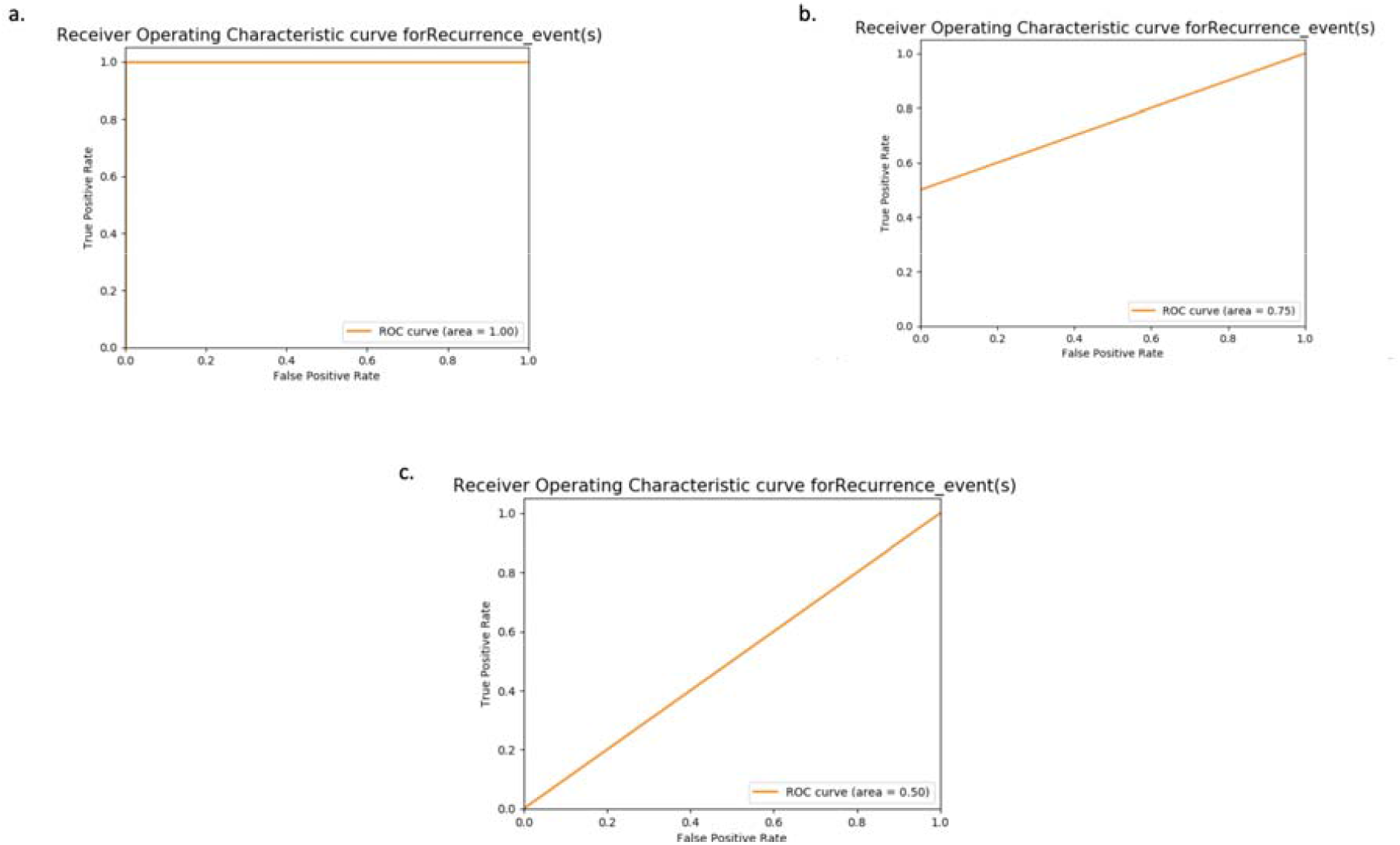
AUCs for prediction of distant recurrence flag using clinicopatholofical profiles using Random Forest and Support Vector Classifier models (p<0.002) in ‘a’ and ‘b’ section respectively, and the same for MultiLayer Perceptron Classifier model (in section ‘c’) during the test phase of the model-training round.

**Table 1:** Performance of various classifiers through a test dataset

| Models | AUC | $R^2$ |
| --- | --- | --- |
| Random Forest | 1.0 | 1.0 |
| Support Vector Classifier | 0.75 | 0.43 |
| Multilayer Perceptron Classifier | 0.5 | -0.7 |

Secondly, the model-type was set to C-Support Vector Classification (SVC) and ran with default model parameters: cv=2, test_size=0.1, pre-ML correlation-threshold set to “-1.0”, data-normalization to “Stand Scaler” and label-normalization to “none”. Further, the grid_search was set to ‘True’ to perform grid search for model through polynomial degree hyperparameters, i.e. kernel=[‘poyl’], degree=[3,4,5,6,7,8,9] and cv=2 (Supplementary Report 2, Figure 1b and Table 1).

Lastly, a supervised neural network, aka multi-layer perceptron (MLP) classifier was employed with parameters: a) pre-ML correlation threshold set to “-1.0”, b) test_size to 0.2, c) cv set to 2 and c) data-normalization set to “Stand Scaler”, and label-normalization to “none”. Further, a grid search was executed using the following hyperparameters: solver: {‘sgd’}, learning_rate: {‘invscaling’}, ‘hidden_layer_sizes’: {(5,), (8,), (9,), (10,), (12,), (15,), (25,), (50,), (100,), (150,), (200,)}, power_t: {0.1, 0.2, 0.25, 0.5, 0.55, 0.6, 0.8, 0.9}, alpha: {0.0001, 0.001, 0.01, 0.1}, ‘cv’: 2, ‘learning_rate_init’: {0.001, 0.01, 0.1} (Supplementary Report 3, Figure 1c and Table 1).

### Validation using ImaGene

Further, the validation of the best of the three models was performed using the similar features from the 9 out of 67 patients (filtered for administration of neoadjuvant therapy and follow up data post adjuvant therapy at an external site (i.e. DHMC)). The validation report yielded by ImaGene showcased the performance of the validation dataset through the model (Supplementary Report 4, Table 2 and Figure 2) as also depicted in the Results section.

**Table 2:** Metrics from validation of the Random Forest model on an external patient-cohort

| Model | AUC | R<sup>2</sup> |
| --- | --- | --- |
| Random Forest | 0.75 | 0.33 |

**Figure 2:**
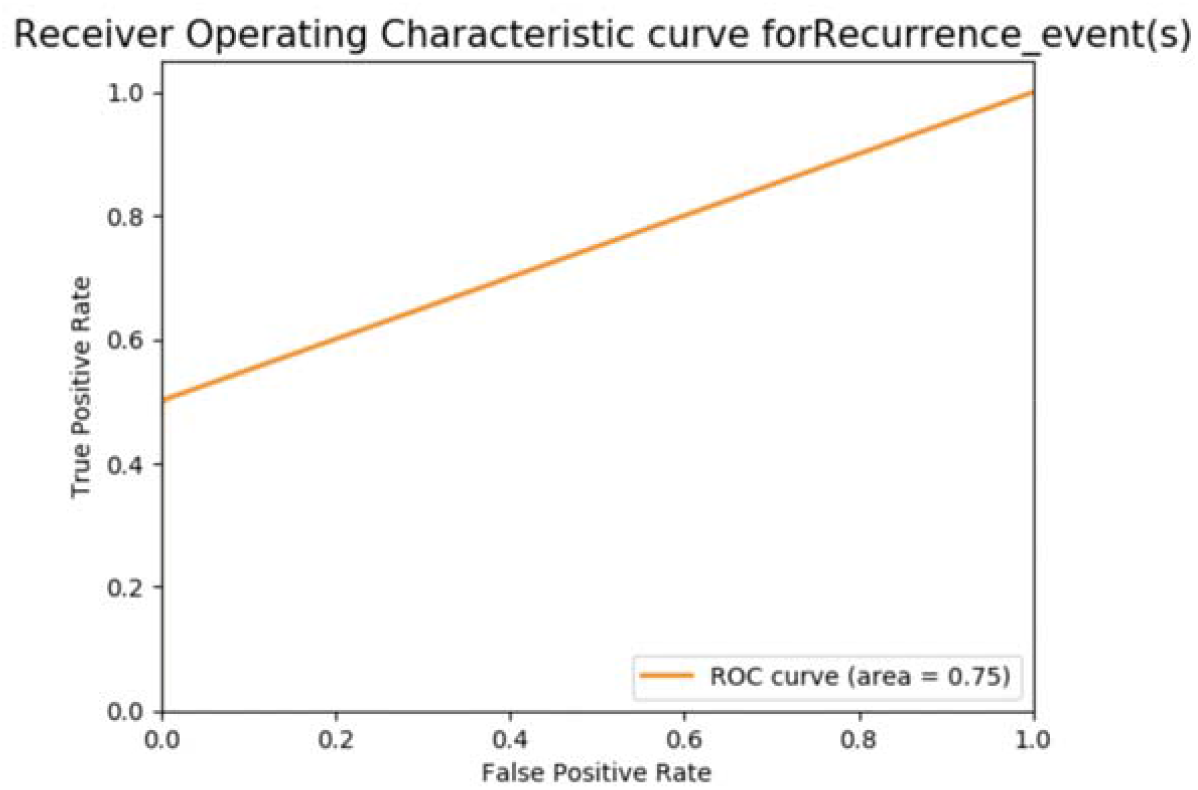
AUC from the validation of Random Forest for the prediction of distant recurrence flags from the clinicopathological profiles in patients from an external site, i.e. DHMC hospital.

## Results

### Model Performance

The runs through ImaGene yielded a detailed performance report along with the supporting tables for the evaluation of each of three models (i.e., Random Forest, SVC and MLP) (Supplementary Report 1-3, Figure 1 and Table 1).

### Random Forest

The parameters set for the training of Random Forest model were as follows: a) “Train” mode with n=161 patients, setting test size to “0.1 (aka 10% of dataset allocated to test)” which partitioned the dataset into train (n_Train_=144) and test (n_Test_=17) sets, b) the K-fold cross validation splitter parameter (i.e., ‘cv’) was set to “2”, and c) Grid search was set to “True” to enable execution of grid search through the hyperparameters: max_depth = [6, 9, 10, 12, 15, 20] and cv=4. The “data” normalization method was set to “StandScaler” while the “label” normalization set to “none”. Further, the pre-ML absolute correlation threshold parameter was set to “-1.0” in order to silent pre-ML filtering and intake all the clinicopathological features for the training of the model. The best score for grid search cross validation (CV) during the training of the model was 0.8, with the Mean Square error (MSE), AUC and R^2^ for the prediction of distant recurrence flags in test dataset (n_test_=17) reported at 0.0, 1.0 and 1.0 respectively at the p-value of less than 0.002 during the permutation test as performed by ImaGene (Sukhadia et al. 2022) (Supplementary Report 1, Figure 1a and Table 1).

### Support Vector Classifier (SVC)

The SVC model ran with the parameters: a) test_size=0.1, b) cv=2, c) pre-ML correlation-threshold set to “-1.0” and d) data-normalization to “Stand Scaler” and label-normalization to “none”, did not yield AUC beyond 0.5. Further, attempting SVC training with GridSearch set to “True” and setting hyperparameters to: a) kernel=[‘poyl’], b) degree=[3,4,5,6,7,8,9] and c) cv=2 yielded the AUC of 0.75 (p<0.002), MSE of 0.06 and R^2^ of 0.43 for the test dataset with the best grid search CV score of 0.78 achieved during the training of the model (Supplementary Report 2, Figure 1b and Table 1).

### Multilayer Perceptron Classifier (MLP)

MLP ran with parameters: a) pre-ML correlation threshold set to “-1.0”, b) test_size to 0.1, c) cv set to 2 and c) data-normalization set to “Stand Scaler”, and label-normalization to “none”, did not yield AUC beyond 0.46 and R^2^ beyond 0. Further, a grid search was executed using the following hyperparameters: solver: {‘sgd’}, learning_rate: {‘invscaling’}, ‘hidden_layer_sizes’: {(5,), (8,), (9,), (10,), (12,), (15,), (25,), (50,), (100,), (150,), (200,)}, power_t: {0.1, 0.2, 0.25, 0.5, 0.55, 0.6, 0.8, 0.9}, alpha: {0.0001, 0.001, 0.01, 0.1}, ‘cv’: 2, ‘learning_rate_init’: {0.001, 0.01, 0.1} did not yield AUC beyond 0.5 and R^2^ beyond 0 (Supplementary Report 3, Figure 1c and Table 1).

The results from several models above alluded Random Forest to be the best performing model for the prediction of distant recurrence flags in IBC patients using their clinicopathological profile (Supplementary Reports 1-3, Figure 1 and Table 2). Therefore, Random Forest model chosen to be validated next on the external patient-cohort, i.e., from DHMC hospital (Supplementary Report 4, Figure 2 and Table 2)

### Model Validation

Further, the validation of the Random Forest model was performed using exact features from n_Validate_=9 out of 67 patients at DHMC (i.e., an external site) which yielded the MSE of 0.125 (i.e., RMSE of 0.35), AUC of 0.75 and R^2^ of 0.33 (Supplementary Report 4, Table 2 and Figure 2), thereby depicting a considerable performance for the external cohort of IBC patient. Augmenting the validation pool with more patients through collaboration with multiple hospitals and universities across the globe through a robust ML/AI platform such as ImaGene could allow us validate the model better in the near future.

## Discussion

We set out to determine the best performing model to predict recurrences in breast cancer patients using clinicopathological measurements of their tumor at diagnostic and therapeutical time-points. These measurements from 161 breast cancer patients from Duke University and nine patients from DHMC included clinical staging of the primary tumor and lymph nodes before neoadjuvant chemotherapy, response to neoadjuvant therapy assessed clinically and pathologically, and administration of adjuvant chemotherapy or anti-*HER2/neu* therapy. These predictors have been used by clinicians previously to predict recurrences in breast cancer (Yau et al. 2022). We used this data to train and test a variety of ML/AI models such as Random Forest, SVC and MLP with 90:10 (train:test) ratio to predict disease recurrence using a robust ML/AI platform: ImaGene (Sukhadia et al. 2022). Our analysis revealed several scores that allowed us to determine the best performing model. We found that the Random Forest model was the best performing model which reported an AUC and R^2^ of 1.0 (p<0.002) for the test dataset compared to other two models(i.e., SVC and MLP). Though SVC reported a considerable performance (AUC=0.75 and R^2^=0.43), MLP couldn’t perform beyond AUC of 0.5 and R^2^ of 0.

Further, the validation of the Random Forest model with the external validation set, i.e., n_validate=_9 patients, showed a considerable AUC of 0.75. One limitation to this study is the low number of samples in the validation set. Our next step to enhance the validation is to expand by collating data from multiple institutions. This could be done using an open-source platform such as ImaGene that enables the democratization of such multi-omic analyses and open-access of the results (Sukhadia et al. 2022).

Our study is the first of its kind to predict distant recurrence in invasive breast cancer patients using clinicopathologic data that includes both pathology and imaging features as well as therapy outcomes across two institutions. The best performing model was the Random Forest, which needs to be validated further in larger multi-institutional studies to encompass a wide range of patient demographics. This model predicts the possibility of distant recurrences in patients, which could provide clinicians with timely information to foresee such recurrences and tailor treatment and management plans accordingly. One of the limitations of this study is the lack of information regarding the distant recurrence site, which is not provided by the dataset shared by Duke University researchers at TCIA platform. Given that additional information, this study could be further extended to potentially predict the site of distant recurrence.

Further, our study supports the capability of ImaGene to utilize several multiomic features of a patient’s tumor throughout the patient’s diagnostic and therapeutic journey. Using Imagene, a patient’s unique pathologic, radiologic, and therapeutic information can be leveraged together to predict the distant recurrences in invasive breast carcinoma. This same model could potentially be utilized to predict disease progression for other cancer types in the future. Our study advances the field of non-invasive prediction of cancer metastasis, and further research in this field may aid researchers and clinicians in identifying disease recurrence, optimizing cancer treatment, and reducing cancer mortality.

## Supporting information

Supplementary Report 1

Supplementary Report 2

Supplementary Report 3

Supplementary Report 4

