## Supplementary Report 1 for "Predicting Distant Recurrences in Invasive Breast Carcinoma Patients Using Clinicopathological Data: A cross-institutional and AI-based study": Supplementary_Report_1_.html

### Radiogenomics Analysis Report

##### 30/12/2022 15:30:54

#### ----------------------------Model inputs-------------------------------

##### Mode:Train

##### Model:RandomForest

##### Params:default

##### Grid\_Params:{'max\_depth': [6, 9, 10, 12, 15, 20], 'cv': 4}

##### No. of imaging features provided: 8

##### No. of gene features provided:1

##### SampleID check results: 'The SampleIDs match for imaging and gene features'

##### No. of samples: 161

#### --------------------------Multivariate Correlations (pearson based)-----------------------


### ----------------------------------Features with highly significant correlations-------------------------------------

#### Below is the list of imaging features

###### ['Adjuvant\_Anti\_Her2\_Neu\_Therapy ', 'Adjuvant\_Chemotherapy', 'Clinical\_Response\_Evaluated\_Through\_Imaging', 'Pathologic\_Response\_to\_Neoadjuvant\_Therapy', 'Pathologic\_response\_to\_Neoadjuvant\_therapy\_Path\_stage\_N', 'Pathologic\_response\_to\_Neoadjuvant\_therapy\_Path\_stage\_T', 'Staging\_Nodes', 'Staging\_Tumor\_size']

#### Below is the list of gene features

###### ['Recurrence\_event(s)']

#### Number of imaging features

### 8

#### Number of gene features

### 1

##### performing Stand\_scaler normalization for imaging features for TRAIN set

##### performing Stand\_scaler normalization for imaging features for TEST set

### -------------------------------Number of Samples for Training and Testing---------------------------------

##### No. of samples for training:144

##### No. of samples for test:17

#### --------------------------Model Summary-----------------------

##### Model Type : RandomForest

###### Grid Search Metrics

###### Best Score : 0.7986111111111112

##### Model Parameters:

###### n\_jobs:None

###### verbose:0

###### estimator\_\_min\_impurity\_decrease:0.0

###### estimator\_\_max\_features:auto

###### estimator\_\_max\_depth:None

###### param\_grid:{'max\_depth': [6, 9, 10, 12, 15, 20]}

#### cv:4

###### scoring:None

###### estimator\_\_criterion:gini

###### estimator\_\_max\_leaf\_nodes:None

###### estimator\_\_verbose:0

###### estimator\_\_n\_jobs:None

###### estimator\_\_min\_samples\_leaf:1

###### estimator\_\_oob\_score:False

###### fit\_params:None

###### estimator\_\_min\_samples\_split:2

###### estimator\_\_warm\_start:False

###### refit:True

###### iid:warn

###### estimator\_\_bootstrap:True

###### estimator\_\_min\_weight\_fraction\_leaf:0.0

###### estimator\_\_n\_estimators:warn

###### pre\_dispatch:2\*n\_jobs

###### estimator\_\_min\_impurity\_split:None

###### estimator\_\_class\_weight:None

###### estimator\_\_random\_state:None

###### return\_train\_score:warn

###### estimator:RandomForestClassifier(bootstrap=True, class\_weight=None, criterion='gini', max\_depth=None, max\_features='auto', max\_leaf\_nodes=None, min\_impurity\_decrease=0.0, min\_impurity\_split=None, min\_samples\_leaf=1, min\_samples\_split=2, min\_weight\_fraction\_leaf=0.0, n\_estimators='warn', n\_jobs=None, oob\_score=False, random\_state=None, verbose=0, warm\_start=False)

###### error\_score:raise-deprecating

#### ----------------Model evaluation for Train data--------------------

##### Min Square Error for the Model

###### MSE of train\_eval set:0.0625

###### No. of features showing LOW 'RMSE/Stdev' (<=1.0): 1

###### All such features with their Low 'RMSE/Stdev' values could be found in output file: train\_eval\_RandomForest\_Labels\_with\_Low\_Ratio.csv No. of features showing HIGH 'RMSE/Stdev' (>1.0): 0 All such features with their High 'RMSE/Stdev' values could be found in output file: train\_eval\_RandomForest\_Labels\_with\_High\_Ratio.csv Model evaluation for Train data for label features showing Low 'RMSE/Stdev' (<=1.0) Content-type: text/html Content-type: text/html ----------------Model evaluation for Test data-------------------- Min Square Error for the Model MSE of test\_eval set:0.0 No. of features showing LOW 'RMSE/Stdev' (<=1.0): 1 All such features with their Low 'RMSE/Stdev' values could be found in output file: test\_eval\_RandomForest\_Labels\_with\_Low\_Ratio.csv No. of features showing HIGH 'RMSE/Stdev' (>1.0): 0 All such features with their High 'RMSE/Stdev' values could be found in output file: test\_eval\_RandomForest\_Labels\_with\_High\_Ratio.csv Model evaluation for Test data for label features showing Low 'RMSE/Stdev' (<=1.0) Content-type: text/html Content-type: text/html ----------------Model evaluation for Validation data-------------------- Min Square Error for the Model MSE of validation\_permut\_1\_Recurrence\_event(s) set:0.117647058824 No. of features showing LOW 'RMSE/Stdev' (<=1.0): 0 All such features with their Low 'RMSE/Stdev' values could be found in output file: validation\_permut\_1\_Recurrence\_event(s)\_RandomForest\_Labels\_with\_Low\_Ratio.csv No. of features showing HIGH 'RMSE/Stdev' (>1.0): 1 All such features with their High 'RMSE/Stdev' values could be found in output file: validation\_permut\_1\_Recurrence\_event(s)\_RandomForest\_Labels\_with\_High\_Ratio.csv Model evaluation for Validation data for label features showing Low 'RMSE/Stdev' (<=1.0) Content-type: text/html Content-type: text/html ----------------Model evaluation for Validation data-------------------- Min Square Error for the Model MSE of validation\_permut\_2\_Recurrence\_event(s) set:0.117647058824 No. of features showing LOW 'RMSE/Stdev' (<=1.0): 0 All such features with their Low 'RMSE/Stdev' values could be found in output file: validation\_permut\_2\_Recurrence\_event(s)\_RandomForest\_Labels\_with\_Low\_Ratio.csv No. of features showing HIGH 'RMSE/Stdev' (>1.0): 1 All such features with their High 'RMSE/Stdev' values could be found in output file: validation\_permut\_2\_Recurrence\_event(s)\_RandomForest\_Labels\_with\_High\_Ratio.csv Model evaluation for Validation data for label features showing Low 'RMSE/Stdev' (<=1.0) Content-type: text/html Content-type: text/html ----------------Model evaluation for Validation data-------------------- Min Square Error for the Model MSE of validation\_permut\_3\_Recurrence\_event(s) set:0.117647058824 No. of features showing LOW 'RMSE/Stdev' (<=1.0): 0 All such features with their Low 'RMSE/Stdev' values could be found in output file: validation\_permut\_3\_Recurrence\_event(s)\_RandomForest\_Labels\_with\_Low\_Ratio.csv No. of features showing HIGH 'RMSE/Stdev' (>1.0): 1 All such features with their High 'RMSE/Stdev' values could be found in output file: validation\_permut\_3\_Recurrence\_event(s)\_RandomForest\_Labels\_with\_High\_Ratio.csv Model evaluation for Validation data for label features showing Low 'RMSE/Stdev' (<=1.0) Content-type: text/html Content-type: text/html ----------------Model evaluation for Validation data-------------------- Min Square Error for the Model MSE of validation\_permut\_4\_Recurrence\_event(s) set:0.0 No. of features showing LOW 'RMSE/Stdev' (<=1.0): 1 All such features with their Low 'RMSE/Stdev' values could be found in output file: validation\_permut\_4\_Recurrence\_event(s)\_RandomForest\_Labels\_with\_Low\_Ratio.csv No. of features showing HIGH 'RMSE/Stdev' (>1.0): 0 All such features with their High 'RMSE/Stdev' values could be found in output file: validation\_permut\_4\_Recurrence\_event(s)\_RandomForest\_Labels\_with\_High\_Ratio.csv Model evaluation for Validation data for label features showing Low 'RMSE/Stdev' (<=1.0) Content-type: text/html Content-type: text/html ----------------Model evaluation for Validation data-------------------- Min Square Error for the Model MSE of validation\_permut\_5\_Recurrence\_event(s) set:0.117647058824 No. of features showing LOW 'RMSE/Stdev' (<=1.0): 0 All such features with their Low 'RMSE/Stdev' values could be found in output file: validation\_permut\_5\_Recurrence\_event(s)\_RandomForest\_Labels\_with\_Low\_Ratio.csv No. of features showing HIGH 'RMSE/Stdev' (>1.0): 1 All such features with their High 'RMSE/Stdev' values could be found in output file: validation\_permut\_5\_Recurrence\_event(s)\_RandomForest\_Labels\_with\_High\_Ratio.csv Model evaluation for Validation data for label features showing Low 'RMSE/Stdev' (<=1.0) Content-type: text/html Content-type: text/html ----------------Model evaluation for Validation data-------------------- Min Square Error for the Model MSE of validation\_permut\_6\_Recurrence\_event(s) set:0.117647058824 No. of features showing LOW 'RMSE/Stdev' (<=1.0): 0 All such features with their Low 'RMSE/Stdev' values could be found in output file: validation\_permut\_6\_Recurrence\_event(s)\_RandomForest\_Labels\_with\_Low\_Ratio.csv No. of features showing HIGH 'RMSE/Stdev' (>1.0): 1 All such features with their High 'RMSE/Stdev' values could be found in output file: validation\_permut\_6\_Recurrence\_event(s)\_RandomForest\_Labels\_with\_High\_Ratio.csv Model evaluation for Validation data for label features showing Low 'RMSE/Stdev' (<=1.0) Content-type: text/html Content-type: text/html ----------------Model evaluation for Validation data-------------------- Min Square Error for the Model MSE of validation\_permut\_7\_Recurrence\_event(s) set:0.117647058824 No. of features showing LOW 'RMSE/Stdev' (<=1.0): 0 All such features with their Low 'RMSE/Stdev' values could be found in output file: validation\_permut\_7\_Recurrence\_event(s)\_RandomForest\_Labels\_with\_Low\_Ratio.csv No. of features showing HIGH 'RMSE/Stdev' (>1.0): 1 All such features with their High 'RMSE/Stdev' values could be found in output file: validation\_permut\_7\_Recurrence\_event(s)\_RandomForest\_Labels\_with\_High\_Ratio.csv Model evaluation for Validation data for label features showing Low 'RMSE/Stdev' (<=1.0) Content-type: text/html Content-type: text/html ----------------Model evaluation for Validation data-------------------- Min Square Error for the Model MSE of validation\_permut\_8\_Recurrence\_event(s) set:0.117647058824 No. of features showing LOW 'RMSE/Stdev' (<=1.0): 0 All such features with their Low 'RMSE/Stdev' values could be found in output file: validation\_permut\_8\_Recurrence\_event(s)\_RandomForest\_Labels\_with\_Low\_Ratio.csv No. of features showing HIGH 'RMSE/Stdev' (>1.0): 1 All such features with their High 'RMSE/Stdev' values could be found in output file: validation\_permut\_8\_Recurrence\_event(s)\_RandomForest\_Labels\_with\_High\_Ratio.csv Model evaluation for Validation data for label features showing Low 'RMSE/Stdev' (<=1.0) Content-type: text/html Content-type: text/html ----------------Model evaluation for Validation data-------------------- Min Square Error for the Model MSE of validation\_permut\_9\_Recurrence\_event(s) set:0.117647058824 No. of features showing LOW 'RMSE/Stdev' (<=1.0): 0 All such features with their Low 'RMSE/Stdev' values could be found in output file: validation\_permut\_9\_Recurrence\_event(s)\_RandomForest\_Labels\_with\_Low\_Ratio.csv No. of features showing HIGH 'RMSE/Stdev' (>1.0): 1 All such features with their High 'RMSE/Stdev' values could be found in output file: validation\_permut\_9\_Recurrence\_event(s)\_RandomForest\_Labels\_with\_High\_Ratio.csv Model evaluation for Validation data for label features showing Low 'RMSE/Stdev' (<=1.0) Content-type: text/html Content-type: text/html ----------------Model evaluation for Validation data-------------------- Min Square Error for the Model MSE of validation\_permut\_10\_Recurrence\_event(s) set:0.117647058824 No. of features showing LOW 'RMSE/Stdev' (<=1.0): 0 All such features with their Low 'RMSE/Stdev' values could be found in output file: validation\_permut\_10\_Recurrence\_event(s)\_RandomForest\_Labels\_with\_Low\_Ratio.csv No. of features showing HIGH 'RMSE/Stdev' (>1.0): 1 All such features with their High 'RMSE/Stdev' values could be found in output file: validation\_permut\_10\_Recurrence\_event(s)\_RandomForest\_Labels\_with\_High\_Ratio.csv Model evaluation for Validation data for label features showing Low 'RMSE/Stdev' (<=1.0) Content-type: text/html Content-type: text/html ----------------Model evaluation for Validation data-------------------- Min Square Error for the Model MSE of validation\_permut\_11\_Recurrence\_event(s) set:0.117647058824 No. of features showing LOW 'RMSE/Stdev' (<=1.0): 0 All such features with their Low 'RMSE/Stdev' values could be found in output file: validation\_permut\_11\_Recurrence\_event(s)\_RandomForest\_Labels\_with\_Low\_Ratio.csv No. of features showing HIGH 'RMSE/Stdev' (>1.0): 1 All such features with their High 'RMSE/Stdev' values could be found in output file: validation\_permut\_11\_Recurrence\_event(s)\_RandomForest\_Labels\_with\_High\_Ratio.csv Model evaluation for Validation data for label features showing Low 'RMSE/Stdev' (<=1.0) Content-type: text/html Content-type: text/html ----------------Model evaluation for Validation data-------------------- Min Square Error for the Model MSE of validation\_permut\_12\_Recurrence\_event(s) set:0.117647058824 No. of features showing LOW 'RMSE/Stdev' (<=1.0): 0 All such features with their Low 'RMSE/Stdev' values could be found in output file: validation\_permut\_12\_Recurrence\_event(s)\_RandomForest\_Labels\_with\_Low\_Ratio.csv No. of features showing HIGH 'RMSE/Stdev' (>1.0): 1 All such features with their High 'RMSE/Stdev' values could be found in output file: validation\_permut\_12\_Recurrence\_event(s)\_RandomForest\_Labels\_with\_High\_Ratio.csv Model evaluation for Validation data for label features showing Low 'RMSE/Stdev' (<=1.0) Content-type: text/html Content-type: text/html ----------------Model evaluation for Validation data-------------------- Min Square Error for the Model MSE of validation\_permut\_13\_Recurrence\_event(s) set:0.117647058824 No. of features showing LOW 'RMSE/Stdev' (<=1.0): 0 All such features with their Low 'RMSE/Stdev' values could be found in output file: validation\_permut\_13\_Recurrence\_event(s)\_RandomForest\_Labels\_with\_Low\_Ratio.csv No. of features showing HIGH 'RMSE/Stdev' (>1.0): 1 All such features with their High 'RMSE/Stdev' values could be found in output file: validation\_permut\_13\_Recurrence\_event(s)\_RandomForest\_Labels\_with\_High\_Ratio.csv Model evaluation for Validation data for label features showing Low 'RMSE/Stdev' (<=1.0) Content-type: text/html Content-type: text/html ----------------Model evaluation for Validation data-------------------- Min Square Error for the Model MSE of validation\_permut\_14\_Recurrence\_event(s) set:0.117647058824 No. of features showing LOW 'RMSE/Stdev' (<=1.0): 0 All such features with their Low 'RMSE/Stdev' values could be found in output file: validation\_permut\_14\_Recurrence\_event(s)\_RandomForest\_Labels\_with\_Low\_Ratio.csv No. of features showing HIGH 'RMSE/Stdev' (>1.0): 1 All such features with their High 'RMSE/Stdev' values could be found in output file: validation\_permut\_14\_Recurrence\_event(s)\_RandomForest\_Labels\_with\_High\_Ratio.csv Model evaluation for Validation data for label features showing Low 'RMSE/Stdev' (<=1.0) Content-type: text/html Content-type: text/html ----------------Model evaluation for Validation data-------------------- Min Square Error for the Model MSE of validation\_permut\_15\_Recurrence\_event(s) set:0.117647058824 No. of features showing LOW 'RMSE/Stdev' (<=1.0): 0 All such features with their Low 'RMSE/Stdev' values could be found in output file: validation\_permut\_15\_Recurrence\_event(s)\_RandomForest\_Labels\_with\_Low\_Ratio.csv No. of features showing HIGH 'RMSE/Stdev' (>1.0): 1 All such features with their High 'RMSE/Stdev' values could be found in output file: validation\_permut\_15\_Recurrence\_event(s)\_RandomForest\_Labels\_with\_High\_Ratio.csv Model evaluation for Validation data for label features showing Low 'RMSE/Stdev' (<=1.0) Content-type: text/html Content-type: text/html ----------------Model evaluation for Validation data-------------------- Min Square Error for the Model MSE of validation\_permut\_16\_Recurrence\_event(s) set:0.117647058824 No. of features showing LOW 'RMSE/Stdev' (<=1.0): 0 All such features with their Low 'RMSE/Stdev' values could be found in output file: validation\_permut\_16\_Recurrence\_event(s)\_RandomForest\_Labels\_with\_Low\_Ratio.csv No. of features showing HIGH 'RMSE/Stdev' (>1.0): 1 All such features with their High 'RMSE/Stdev' values could be found in output file: validation\_permut\_16\_Recurrence\_event(s)\_RandomForest\_Labels\_with\_High\_Ratio.csv Model evaluation for Validation data for label features showing Low 'RMSE/Stdev' (<=1.0) Content-type: text/html Content-type: text/html ----------------Model evaluation for Validation data-------------------- Min Square Error for the Model MSE of validation\_permut\_17\_Recurrence\_event(s) set:0.117647058824 No. of features showing LOW 'RMSE/Stdev' (<=1.0): 0 All such features with their Low 'RMSE/Stdev' values could be found in output file: validation\_permut\_17\_Recurrence\_event(s)\_RandomForest\_Labels\_with\_Low\_Ratio.csv No. of features showing HIGH 'RMSE/Stdev' (>1.0): 1 All such features with their High 'RMSE/Stdev' values could be found in output file: validation\_permut\_17\_Recurrence\_event(s)\_RandomForest\_Labels\_with\_High\_Ratio.csv Model evaluation for Validation data for label features showing Low 'RMSE/Stdev' (<=1.0) Content-type: text/html Content-type: text/html ----------------Model evaluation for Validation data-------------------- Min Square Error for the Model MSE of validation\_permut\_18\_Recurrence\_event(s) set:0.117647058824 No. of features showing LOW 'RMSE/Stdev' (<=1.0): 0 All such features with their Low 'RMSE/Stdev' values could be found in output file: validation\_permut\_18\_Recurrence\_event(s)\_RandomForest\_Labels\_with\_Low\_Ratio.csv No. of features showing HIGH 'RMSE/Stdev' (>1.0): 1 All such features with their High 'RMSE/Stdev' values could be found in output file: validation\_permut\_18\_Recurrence\_event(s)\_RandomForest\_Labels\_with\_High\_Ratio.csv Model evaluation for Validation data for label features showing Low 'RMSE/Stdev' (<=1.0) Content-type: text/html Content-type: text/html ----------------Model evaluation for Validation data-------------------- Min Square Error for the Model MSE of validation\_permut\_19\_Recurrence\_event(s) set:0.117647058824 No. of features showing LOW 'RMSE/Stdev' (<=1.0): 0 All such features with their Low 'RMSE/Stdev' values could be found in output file: validation\_permut\_19\_Recurrence\_event(s)\_RandomForest\_Labels\_with\_Low\_Ratio.csv No. of features showing HIGH 'RMSE/Stdev' (>1.0): 1 All such features with their High 'RMSE/Stdev' values could be found in output file: validation\_permut\_19\_Recurrence\_event(s)\_RandomForest\_Labels\_with\_High\_Ratio.csv Model evaluation for Validation data for label features showing Low 'RMSE/Stdev' (<=1.0) Content-type: text/html Content-type: text/html ----------------Model evaluation for Validation data-------------------- Min Square Error for the Model MSE of validation\_permut\_20\_Recurrence\_event(s) set:0.117647058824 No. of features showing LOW 'RMSE/Stdev' (<=1.0): 0 All such features with their Low 'RMSE/Stdev' values could be found in output file: validation\_permut\_20\_Recurrence\_event(s)\_RandomForest\_Labels\_with\_Low\_Ratio.csv No. of features showing HIGH 'RMSE/Stdev' (>1.0): 1 All such features with their High 'RMSE/Stdev' values could be found in output file: validation\_permut\_20\_Recurrence\_event(s)\_RandomForest\_Labels\_with\_High\_Ratio.csv Model evaluation for Validation data for label features showing Low 'RMSE/Stdev' (<=1.0) Content-type: text/html Content-type: text/html
