## Supplementary Report 2 for "Predicting Distant Recurrences in Invasive Breast Carcinoma Patients Using Clinicopathological Data: A cross-institutional and AI-based study": Supplementary_Report_2_.html

### Radiogenomics Analysis Report

##### 16/04/2023 11:45:32

#### ----------------------------Model inputs-------------------------------

##### Mode:Train

##### Model:SVC

##### Params:default

##### Grid\_Params:{'kernel': ['poly'], 'cv': 2, 'degree': [3, 4, 5, 6, 7, 8, 9]}

##### No. of imaging features provided: 8

##### No. of gene features provided:1

##### SampleID check results: 'The SampleIDs match for imaging and gene features'

##### No. of samples: 161

#### --------------------------Multivariate Correlations (pearson based)-----------------------


### ----------------------------------Features with highly significant correlations-------------------------------------

#### Below is the list of imaging features

###### ['Adjuvant\_Anti\_Her2\_Neu\_Therapy ', 'Adjuvant\_Chemotherapy', 'Clinical\_Response\_Evaluated\_Through\_Imaging', 'Pathologic\_Response\_to\_Neoadjuvant\_Therapy', 'Pathologic\_response\_to\_Neoadjuvant\_therapy\_Path\_stage\_N', 'Pathologic\_response\_to\_Neoadjuvant\_therapy\_Path\_stage\_T', 'Staging\_Nodes', 'Staging\_Tumor\_size']

#### Below is the list of gene features

###### ['Recurrence\_event(s)']

#### Number of imaging features

### 8

#### Number of gene features

### 1

##### performing Stand\_scaler normalization for imaging features for TRAIN set

##### performing Stand\_scaler normalization for imaging features for TEST set

### -------------------------------Number of Samples for Training and Testing---------------------------------

##### No. of samples for training:144

##### No. of samples for test:17

#### --------------------------Model Summary-----------------------

##### Model Type : SVC

###### Grid Search Metrics

###### Best Score : 0.7847222222222222

##### Model Parameters:

###### n\_jobs:None

###### verbose:0

###### estimator\_\_gamma:auto\_deprecated

###### estimator\_\_decision\_function\_shape:ovr

###### estimator\_\_probability:False

###### param\_grid:{'kernel': ['poly'], 'degree': [3, 4, 5, 6, 7, 8, 9]}

#### cv:2

###### scoring:None

###### estimator\_\_cache\_size:200

###### estimator\_\_verbose:False

###### pre\_dispatch:2\*n\_jobs

###### estimator\_\_kernel:rbf

###### fit\_params:None

###### estimator\_\_max\_iter:-1

###### refit:True

###### iid:warn

###### estimator\_\_shrinking:True

###### estimator\_\_degree:3

###### estimator\_\_class\_weight:None

###### estimator\_\_C:1.0

###### estimator\_\_random\_state:None

###### return\_train\_score:warn

###### estimator:SVC(C=1.0, cache\_size=200, class\_weight=None, coef0=0.0, decision\_function\_shape='ovr', degree=3, gamma='auto\_deprecated', kernel='rbf', max\_iter=-1, probability=False, random\_state=None, shrinking=True, tol=0.001, verbose=False)

###### estimator\_\_coef0:0.0

###### error\_score:raise-deprecating

###### estimator\_\_tol:0.001

#### ----------------Model evaluation for Train data--------------------

##### Min Square Error for the Model

###### MSE of train\_eval set:0.131944444444

###### No. of features showing LOW 'RMSE/Stdev' (<=1.0): 1
