## Supplementary Report 3 for "Predicting Distant Recurrences in Invasive Breast Carcinoma Patients Using Clinicopathological Data: A cross-institutional and AI-based study": Supplementary_Report_3_.html

### Radiogenomics Analysis Report

##### 16/04/2023 12:56:53

#### ----------------------------Model inputs-------------------------------

##### Mode:Train

##### Model:MLPClassifier

##### Params:default

##### Grid\_Params:{'hidden\_layer\_sizes': [(5,), (8,), (9,), (10,), (12,), (15,), (25,), (50,), (100,), (150,), (200,)], 'learning\_rate': ['invscaling'], 'solver': ['sgd'], 'power\_t': [0.1, 0.2, 0.25, 0.5, 0.55, 0.6, 0.8, 0.9], 'alpha': [0.0001, 0.001, 0.01, 0.1], 'cv': 2, 'learning\_rate\_init': [0.001, 0.01, 0.1]}

##### No. of imaging features provided: 8

##### No. of gene features provided:1

##### SampleID check results: 'The SampleIDs match for imaging and gene features'

##### No. of samples: 161

#### --------------------------Multivariate Correlations (pearson based)-----------------------


### ----------------------------------Features with highly significant correlations-------------------------------------

#### Below is the list of imaging features

###### ['Adjuvant\_Anti\_Her2\_Neu\_Therapy ', 'Adjuvant\_Chemotherapy', 'Clinical\_Response\_Evaluated\_Through\_Imaging', 'Pathologic\_Response\_to\_Neoadjuvant\_Therapy', 'Pathologic\_response\_to\_Neoadjuvant\_therapy\_Path\_stage\_N', 'Pathologic\_response\_to\_Neoadjuvant\_therapy\_Path\_stage\_T', 'Staging\_Nodes', 'Staging\_Tumor\_size']

#### Below is the list of gene features

###### ['Recurrence\_event(s)']

#### Number of imaging features

### 8

#### Number of gene features

### 1

##### performing Stand\_scaler normalization for imaging features for TRAIN set

##### performing Stand\_scaler normalization for imaging features for TEST set

### -------------------------------Number of Samples for Training and Testing---------------------------------

##### No. of samples for training:144

##### No. of samples for test:17

#### --------------------------Model Summary-----------------------

##### Model Type : MLPClassifier

###### Grid Search Metrics

###### Best Score : 0.8402777777777778

##### Model Parameters:

###### estimator\_\_epsilon:1e-08

###### n\_jobs:None

###### estimator\_\_hidden\_layer\_sizes:(100,)

###### verbose:0

###### estimator\_\_early\_stopping:False

###### estimator\_\_nesterovs\_momentum:True

###### estimator\_\_alpha:0.0001

###### param\_grid:{'hidden\_layer\_sizes': [(5,), (8,), (9,), (10,), (12,), (15,), (25,), (50,), (100,), (150,), (200,)], 'learning\_rate': ['invscaling'], 'solver': ['sgd'], 'power\_t': [0.1, 0.2, 0.25, 0.5, 0.55, 0.6, 0.8, 0.9], 'alpha': [0.0001, 0.001, 0.01, 0.1], 'learning\_rate\_init': [0.001, 0.01, 0.1]}

###### estimator\_\_shuffle:True

###### scoring:None

#### cv:2

###### estimator\_\_activation:relu

###### estimator\_\_verbose:False

###### estimator\_\_learning\_rate\_init:0.001

###### fit\_params:None

###### estimator\_\_solver:adam

###### estimator\_\_warm\_start:False

###### estimator\_\_max\_iter:200

###### refit:True

###### estimator\_\_n\_iter\_no\_change:10

###### estimator\_\_beta\_2:0.999

###### estimator\_\_beta\_1:0.9

###### pre\_dispatch:2\*n\_jobs

###### estimator\_\_power\_t:0.5

###### estimator\_\_learning\_rate:constant

###### iid:warn

###### estimator\_\_random\_state:None

###### estimator\_\_batch\_size:auto

###### return\_train\_score:warn

###### estimator\_\_momentum:0.9

###### estimator:MLPClassifier(activation='relu', alpha=0.0001, batch\_size='auto', beta\_1=0.9, beta\_2=0.999, early\_stopping=False, epsilon=1e-08, hidden\_layer\_sizes=(100,), learning\_rate='constant', learning\_rate\_init=0.001, max\_iter=200, momentum=0.9, n\_iter\_no\_change=10, nesterovs\_momentum=True, power\_t=0.5, random\_state=None, shuffle=True, solver='adam', tol=0.0001, validation\_fraction=0.1, verbose=False, warm\_start=False)

###### error\_score:raise-deprecating

###### estimator\_\_validation\_fraction:0.1

###### estimator\_\_tol:0.0001

#### ----------------Model evaluation for Train data--------------------

##### Min Square Error for the Model

###### MSE of train\_eval set:0.180555555556

###### No. of features showing LOW 'RMSE/Stdev' (<=1.0): 0

###### All such features with their Low 'RMSE/Stdev' values could be found in output file: train\_eval\_MLPClassifier\_Labels\_with\_Low\_Ratio.csv No. of features showing HIGH 'RMSE/Stdev' (>1.0): 1 All such features with their High 'RMSE/Stdev' values could be found in output file: train\_eval\_MLPClassifier\_Labels\_with\_High\_Ratio.csv Model evaluation for Train data for label features showing Low 'RMSE/Stdev' (<=1.0) Content-type: text/html Content-type: text/html ----------------Model evaluation for Test data-------------------- Min Square Error for the Model MSE of test\_eval set:0.294117647059 No. of features showing LOW 'RMSE/Stdev' (<=1.0): 0 All such features with their Low 'RMSE/Stdev' values could be found in output file: test\_eval\_MLPClassifier\_Labels\_with\_Low\_Ratio.csv No. of features showing HIGH 'RMSE/Stdev' (>1.0): 1 All such features with their High 'RMSE/Stdev' values could be found in output file: test\_eval\_MLPClassifier\_Labels\_with\_High\_Ratio.csv Model evaluation for Test data for label features showing Low 'RMSE/Stdev' (<=1.0) Content-type: text/html Content-type: text/html
